# Activity-based probes and chemical proteomics uncover the biological impact of targeting HMGCS1

**DOI:** 10.1101/2025.05.21.655359

**Authors:** Sang Ah Yi, Liang Sun, Yi Rao, Alban Ordureau, Jason S. Lewis, Heeseon An

## Abstract

Mevalonate is a precursor for essential metabolites, such as isoprenoids and sterols. Its synthesis starts with HMGCS1 producing HMG-CoA, which is then converted to mevalonate by HMGCR, a target of statins. Cancer cells often upregulate enzymes in the mevalonate pathway (MVP) to meet their metabolic demands, leading to the development of inhibitors targeting several enzymes in this pathway. However, current inhibitors have not yet shown significant anti-cancer activity. While HMGCS1 has unique biochemical properties that distinguish it from other MVP enzymes, the effects of inhibiting HMGCS1 have not been thoroughly investigated. Here, we present a set of chemical probes that enable us to systematically assess the proteome-wide selectivity and potency of Hymeglusin, the primary inhibitor of HMGCS1 used in the field, confirming it as a useful tool for short-term HMGCS1 inhibition. Inhibiting HMGCS1 with Hymeglusin causes proteome changes that are nearly identical to those caused by inhibiting HMGCR or degrading HMGCS1. Accordingly, simultaneously targeting HMGCS1 and HMGCR effectively suppresses the growth of statin-resistant cells and xenograft models, without increasing the risk of side effects. Finally, we find that while Hymeglusin is a valuable tool for short-term mechanistic studies, its usefulness is limited for long-term efficacy studies due to its poor stability in serum. Together, this study highlights the biological implications of targeting HMGCS1 as monotherapy or in combination with statins, and caution is required when using Hymeglusin as a tool.

## INTRODUCTION

Mevalonate is an essential precursor for the biosynthesis of sterols, isoprenoids, vitamin D, and ubiquinone.(1, 2) Consequently, the enzymes involved in the mevalonate pathway (MVP) play crucial roles in multiple aspects of cellular physiology, from membrane biology to cell signaling and mitochondrial metabolism.(3) The synthesis of mevalonate begins with the first step, catalyzed by 3-Hydroxy-3-Methylglutaryl-Coenzyme A Synthase 1 (HMGCS1), which condenses acetyl-CoA with acetoacetyl-CoA to produce HMG-CoA. The HMG-CoA then undergoes reduction to mevalonate via HMG-CoA Reductase (HMGCR). Following this, approximately 20 downstream enzymes convert mevalonate into sterols and other non-sterol end-products. Growing evidence indicates that cancer cells exhibit upregulation of MVP enzymes, which may stimulate glucose uptake and various signaling pathways, including RAS, RAB, and RHO.(4–8) This upregulation has also been linked to resistance to various chemotherapies, emphasizing the significance of these enzymes in cancer progression and their ability to manage stress.(9–11) Prior research suggests that inhibiting the mevalonate pathway induces apoptotic death of cancer cells by blocking the geranylgeranylation of proteins.(12, 13) Consequently, several enzymes in this pathway, such as HMGCR, FDPS, GGDPS1, FTase, and GGTase, have been exploited as potential pharmacological targets.(5, 7) However, the existing approaches or inhibitors have yet to produce significant clinical anti-cancer activity.(7, 14–21)

HMGCS1 represents an underexplored pharmacological target within the mevalonate pathway, despite the fact that it possesses several attractive features that distinguish it from HMGCR and other downstream MVP enzymes. Notably, HMGCS1 contains a catalytic cysteine in its active site that can be readily leveraged for developing covalent inhibitors and chemical probes to enhance potency and sustained efficacy compared to non-covalent inhibitors. Moreover, we previously demonstrated that HMGCS1 levels are directly regulated by the master regulator of cell growth, mTORC1,(22) and an increased requirement for HMGCS1 during the activation of cell proliferation signals makes it a limiting enzyme in cells with hyperactive mTORC1, a frequently observed feature of cancer cells. Thus, perturbation of HMGCS1 can be an alternative strategy to treat cancers exhibiting upregulated metabolites and signaling pathways related to MVP. (22)

However, the chemical tools available to perturb the HMGCS1 activity are currently limited, with Hymeglusin serving as the primary small molecule probe.(10, 23, 24) Identified as an HMGCS1 inhibitor in 1987, Hymeglusin was isolated from Fusarium and Scopulariopsis fungi.(25–30) The co-crystal structure of HMGCS1 and Hymeglusin revealed that Hymeglusin forms a thioester bond with the catalytic cysteine of HMGCS1 following the opening of its lactone ring.(31) A subsequent study showed that this HMGCS1-Hymeglusin adduct is stable *in vitro*, with an estimated half-life of 55 hours.(26) The same study also showed that Hymeglusin-mediated inhibition of HMGCS1 in cells is reversed within 30 minutes after withdrawal when measured by the conversion of radioactive acetate into cholesterol.(26) However, the occupancy level of the intracellular HMGCS1 catalytic cysteine and the cause of this increased reversibility of Hymeglusin remain unclear, primarily due to the lack of tools that allow direct monitoring. Additionally, a comprehensive quantitative analysis of the reactivity of Hymeglusin toward the global proteome, as well as its effects on reshaping the proteome landscape, has not been reported.

In the present study, we have developed activity-based probes for HMGCS1, functionalized with fluorescent dye or biotin handles, that facilitate the investigation of the effects of covalent inhibition of HMGCS1 in human cells. By combining these probes with proteomics analyses, we show that Hymeglusin specifically reacts with the catalytic cysteine of HMGCS1 in cells. Hymeglusin treatment induces global proteomic changes that are nearly identical to those induced by the HMGCR inhibitor, Simvastatin. Our unbiased data support the selective inhibition of mevalonate synthesis by Hymeglusin and reveal non-redundant functions of HMGCS1 and HMGCR outside the mevalonate pathway. Consequently, the concurrent targeting of both enzymes potentiates inhibition of mevalonate synthesis, leading to increased anti-cancer effects in cells and in mouse xenograft models. We also demonstrate that targeted proteolysis of HMGCS1 exhibits a more potent anti-proliferation effect than inhibition by Hymeglusin, due to its poor serum stability. In conclusion, this study presents a comprehensive toolkit for inhibiting and analyzing HMGCS1 activity, providing insights into the cellular and pharmacological effects of this intervention.

## RESULTS

### Hymeglusin-fluorescence conjugates monitor occupancy of the HMGCS1 active site

HMGCS1 is a 57 kDa protein known to form tight homodimers in cells and *in vitro*.(22, 32) A previous co-crystal structure showed that Hymeglusin forms a covalent bond with the catalytic cysteine of HMGCS1, thereby inhibiting its enzymatic activity (**Fig. 1A**).(33) While Hymeglusin has been used as a representative inhibitor of HMGCS1 for biological studies,(10, 23, 24) its proteome-wide selectivity has not been evaluated quantitatively. This lack of comprehensive evaluation creates uncertainties about its biological implications and on-target effects. We, therefore, synthesized fluorescent probes by conjugating the Hymeglusin warhead to two different dyes, fluorescein or tetramethylrhodamine (TMR), through a linker (**Fig. 1B**). The Hymeglusin-fluorecein (HG-FL) probe was then incubated with either wild-type (WT) or a catalytically inactive HMGCS1 harboring a Cys-Ala substitution *in vitro*, after purifying them from HEK293T cells (**Fig. 1C**). In-gel fluorescence analysis showed a signal exclusively in wild-type HMGCS1, indicating that HG-FL binds the catalytic cysteine residue. When recombinant HMGCS1 was pre-treated in vitro with different concentrations of Hymeglusin before adding HG-FL, the fluorescent signal decreased in a dose-dependent manner (**Fig. 1D**). HG-FL binding to HMGCS1 was disrupted in the presence of a reducing agent, dithiothreitol (DTT), which may cleave the thioester bond formed between Cys129 and Hymeglusin (**Fig. 1E**). However, the strong resistance of this interaction under denaturing conditions implied a tight binding of Hymeglusin to the hydrophobic pocket, which hinders the access of DTT for trans thiolation reaction. We subsequently investigated whether HG-fluorescence probes could be used to monitor the occupancy of the catalytic cysteine residue within intracellular HMGCS1 (**Fig. 1F**, upper panel). WT HEK293T cells were treated with Hymeglusin for 2 hours, followed by cell lysis, and incubation with the HG probes. Subsequent in-gel fluorescence analyses revealed a fluorescence band that was reversed upon pre-incubation of cells with Hymeglusin (**Fig. 1F**, bottom left panel). CRISPR-engineered HCT116 cells expressing endogenous HMGCS1-FKBP12^F36V^ or HMGCS1-mEGFP fusion protein showed similar results when probed by either HG-FL or HG-TMR, with the shifts in molecular weight that match with HMGCS1 chimera (**Fig. 1F**, bottom middle/right panels). The same gels were then subjected to immunoblotting for HMGCS1 detection, and the fluorescent bands coincided with the bands detected by the anti-HMGCS1 antibody. These data indicate that HG-fluorescence probes can serve as activity-based probes for HMGCS1.

**FIGURE 1.**
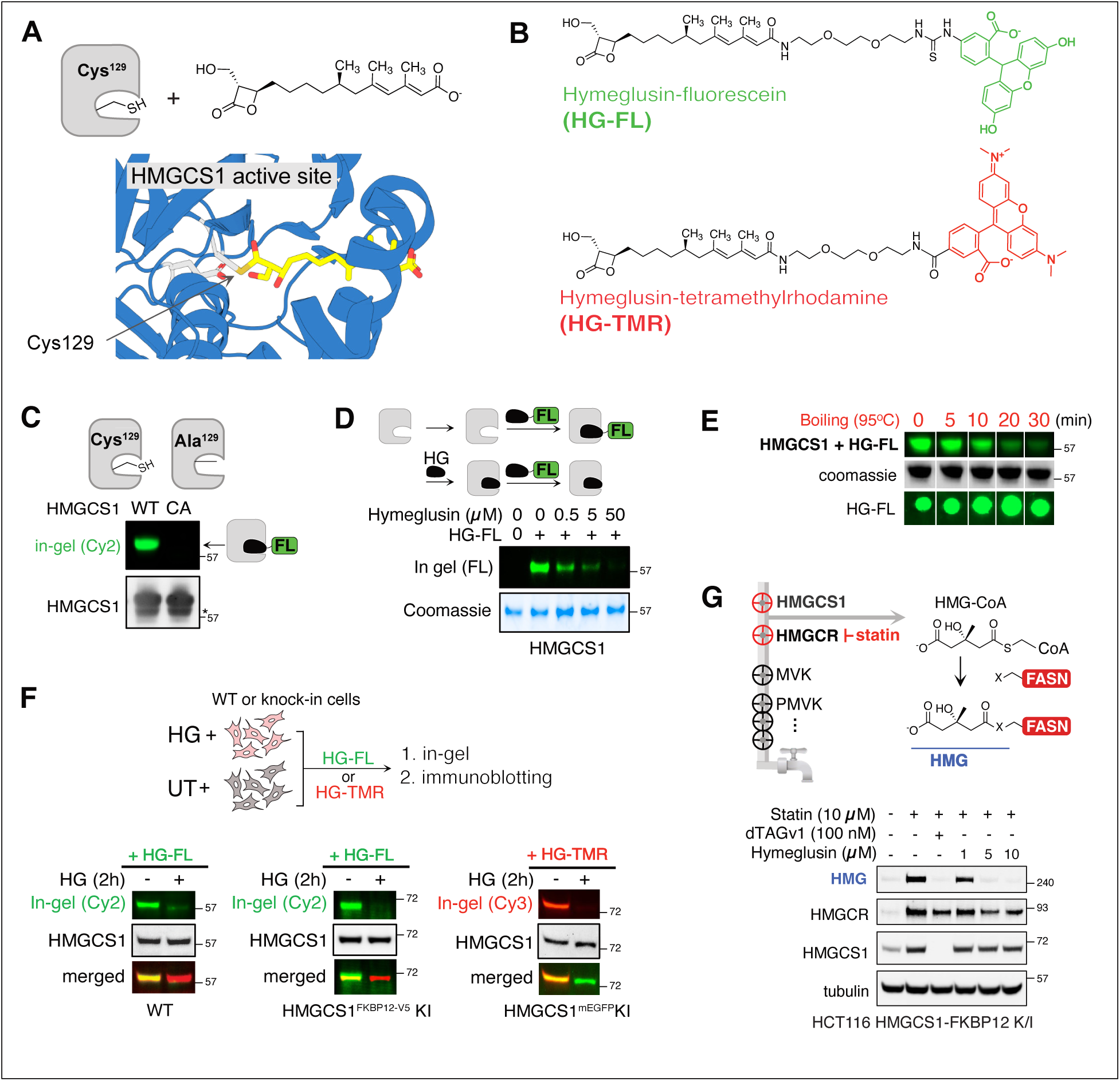
Hymeglusin-fluorescence conjugates serve as activity-based probes for HMGCS1. (A) A schematic of the HMGCS1 and Hymeglusin (top) and the zoomed-in co-crystal structure of the Hymeglusin-bound HMGCS1 (bottom, PDB: 2F9A) (B) Chemical structures of the Hymeglusin-fluorescein (HG-FL) and Hymeglusin-tetramethylrhodamine probes (HG-TMR). (C) Comparative in-gel fluorescence assay of wild-type (WT) or a catalytically inactive HMGCS1 mutant (C129A, CA) confirms the labeling of the catalytic cysteine of HMGCS1 with HG-FL. (D) Top: scheme of the HG-FL competition reaction *in vitro*. Bottom: incubating recombinant HMGCS1 (1 µg) with increasing concentrations of Hymeglusin for 30 minutes results in a decrease of HG-FL labeled HMGCS1. (E) The labeling of HMGCS1 by HG-FL was reversed after boiling at 95°C for at least 20 minutes in a buffer containing 2% LDS and 50 mM DTT. SDS-PAGE resolved HMGCS1 before the in-gel fluorescence analysis, while the stability of the HG-FL probe in the given conditions was assessed by dot blot in parallel. (F) HEK293T WT and HCT116 knock-in (KI) (HMGCS1-FKBP12^F36V^ or HMGCS1-mEGFP) cells were treated with Hymeglusin (4 μM for WT, 0.5 μM for KI, 2 hours) or left untreated, followed by cell lysis and incubation with HG-FL or HG-TMR for one hour. After in-gel fluorescence analysis, the gels were transferred to the PVDF membrane for immunoblotting using an anti-HMGCS1 antibody. (G) HMGCR inhibition by statins leads to the accumulation of HMG-CoA in cells. This results in the non-enzymatic modification of several residues in the FASN active site by the HMG moiety, which can be detected by the anti-HMG antibody (top). HCT116 cells expressing endogenous HMGCS1-FKBP12^F36V^ were treated with the indicated chemicals for 24 hours, followed by immunoblotting analysis using anti-HMG, anti-HMGCR, anti-HMGCS1, and anti-tubulin antibodies (bottom).

Does Hymeglusin occupancy of the HMGCS1 catalytic cysteine lead to the depletion of cellular HMG-CoA, the product of HMGCS1 enzymatic activity? Previous investigations have demonstrated that the accumulation of HMG-CoA in cells, resulting from the inhibition of HMGCR by statins, facilitates a non-enzymatic reaction with fatty acid synthase (FASN), leading to the formation of HMG-modified FASN (**Fig. 1G**, upper panel).(34) The presence of this modification can be effectively monitored using an anti-HMG antibody.(35) Notably, prior studies have shown that the overall reactivity of this antibody correlates with intracellular levels of HMG-CoA. Indeed, when HCT116 cells expressing endogenous HMGCS1-FKBP12^F36V^ fusion protein were treated with Simvastatin, an HMG positive band appeared around 240 kDa range (**Fig. 1G**, bottom panel, first two lanes). The addition of the dTAGv1 ligand, which recruits the VHL-containing E3 ligase complex to the FKBP12^F36V^ tag and induces the degradation of HMGCS1-FKBP12^F36V^chimera, caused the disappearance of the HMGylated protein signal (**Fig. 1G**, bottom panel, third lane, and **Fig. S1A**).(22, 36) This indicates that the cellular HMG-CoA level dropped in the absence of HMGCS1, serving as the positive control for this assay system. Treatment with increasing concentrations of Hymeglusin similarly led to a progressive decrease in the HMGylated protein intensity, indicating inhibition of HMGCS1 activity. Of note, statin treatment induced a substantial increase in HMGCS1 and HMGCR levels due to the feedback effect mediated by the SREBP2 transcription factor.(37)

### Hymeglusin reacts highly selectively with HMGCS1 in human cells

We next evaluated the reactivity of Hymeglusin toward the human proteome using an unbiased chemical proteomics approach. To conclusively identify the Hymeglusin-labeled proteins using mass spectrometry, we synthesized the Hymeglusin-biotin (HG-Biotin) probe (**Fig. 2A**, top). HEK293T cells were either treated with Hymeglusin or DMSO for 30 min prior to incubation with HG-biotin. The proteins labeled with HG-biotin were enriched using streptavidin beads, and the eluates were analyzed using a 6-plex tandem-mass-tag (TMT)-based quantification. Our proteomic analysis revealed that HMGCS1 is the only significantly enriched protein that was displaced by the pretreatment with Hymeglusin (12-fold increase, 50 peptides detected), demonstrating Hymeglusin’s high selectivity for HMGCS1 (**Fig. 2B**). An orthogonal immunoblotting assay confirmed the strong enrichment of HMGCS1 by HG-biotin (**Fig. 2C**). In contrast, endogenously biotinylated proteins such as ACACA (18 peptides), PCCA (19 peptides), and MCCC1 (14 peptides), were enriched regardless of Hymeglusin pre-treatment, as a consequence of their interaction with streptavidin (**Table S1**). Consistently, the HG-FL assay on HEK293T cells that were pretreated with increasing concentrations of Hymeglusin revealed a single strong fluorescence signal around 57 kDa, corresponding to the molecular weight of HMGCS1 (**Fig. 2D**). This signal disappeared when the cells were pretreated with 250-500 nM Hymeglusin. We conclude that Hymeglusin is a potent and selective inhibitor of HMGCS1, making it a suitable tool for examining the downstream biological effects of acute HMGCS1 inhibition.

**FIGURE 2.**
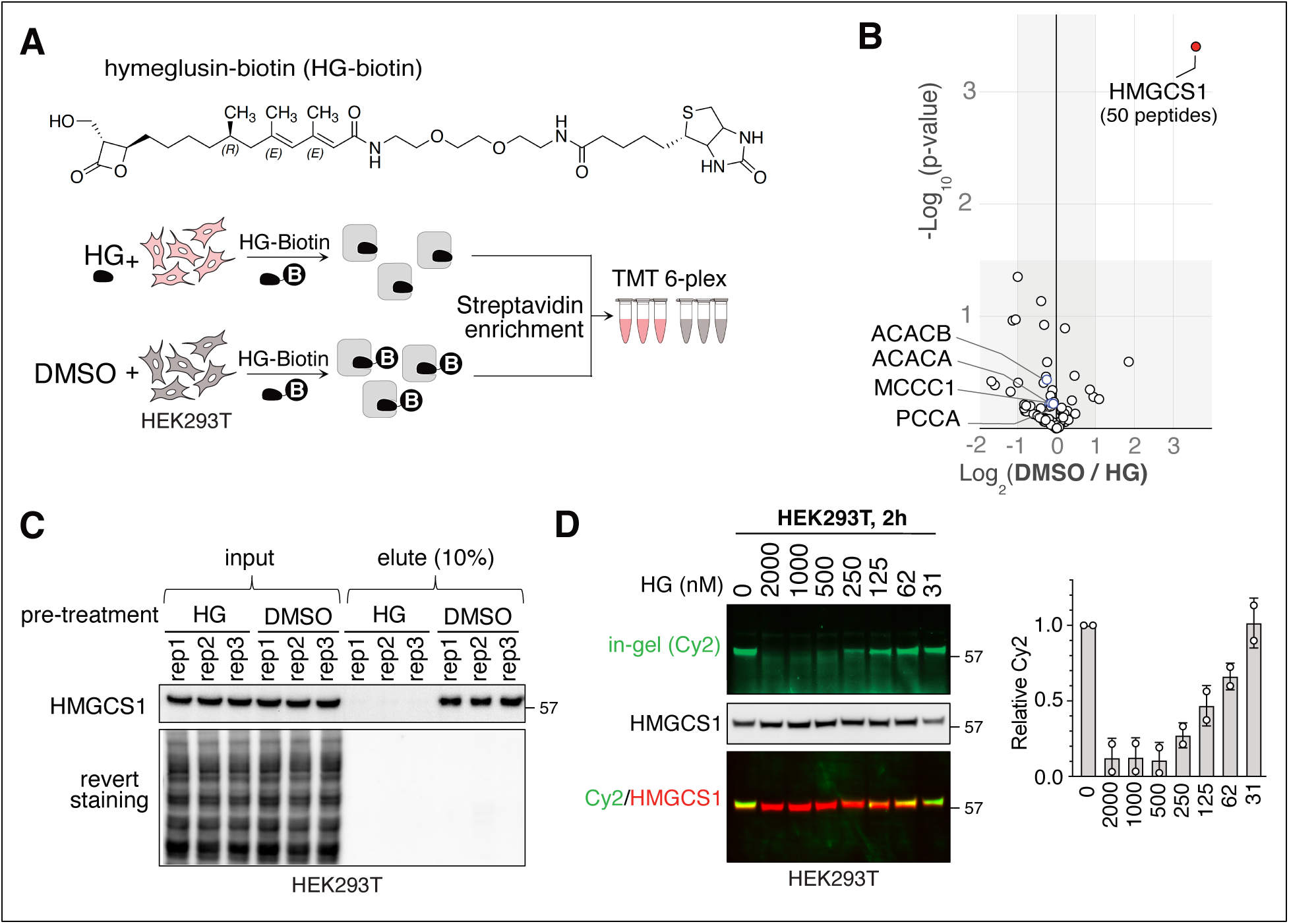
Hymeglusin is a highly selective inhibitor of HMGCS1 in human cells. (A) The structure of the Hymeglusin-biotin probe is shown at the top, and the workflow of competitive affinity purification mass-spectrometry (AP-MS) using Hymeglusin-biotin combined with TMT-6 plex methods is shown at the bottom. (B) HEK293T cells were either treated with HG (1 µM, 30 min) or DMSO, followed by further incubation with the in-house HG-biotin probe (1 µM, 2 h). After streptavidin beads enrichment, the eluates were analyzed through tandem-mass-tag-based proteomic analysis. The proteomic data is presented as a volcano plot of the –log_10_-transformed p-value versus the log_2_-transformed ratio of DMSO/Hymeglusin pre-treated cells. N = 3 biological replicates. p-values were calculated by two-sided Welch’s t-test (adjusted to 1% FDR for multiple comparisons). (C) HEK293T cells were treated as described in panel A. After streptavidin enrichment, the eluates were subjected to immunoblotting using the anti-HMGCS1 antibody. Rep: replicate. (D) HEK293T cells were treated with increasing concentrations of Hymeglusin for 2 hours, followed by HG-FL treatment and in-gel fluorescence analysis. The same extracts were then transferred to a PVDF membrane to probe for HMGCS1 using immunoblotting. The average of two replicates is shown on the right.

### The inhibition or degradation of HMGCS1, or inhibition of HMGCR, results in identical changes to the proteome

To gain a deeper understanding of the cellular response to the inhibition of HMGCS1, we analyzed global proteomic changes following Hymeglusin treatment (**Fig. 3A**, left). We compared the proteome alterations caused by the acute degradation of HMGCS1 by using the HCT116 HMGCS1-FKBP12^F36V^ knock-in cells, which may decouple any side effects of Hymeglusin and non-enzymatic roles of HMGCS1, if they exist. We also compared the proteome changes induced by the inhibition of HMGCR, as HMGCS1 and HMGCR are the first two enzymes in the mevalonate pathway (**Table S2**). If these two enzymes have additional functions outside the mevalonate pathway, it may be evident in the proteome changes. Immunoblotting analysis showed that dTAGv1 treatment for 24 hours led to the expected depletion of HMGCS1 and the upregulation of unprenylated RhoA and HMGCR, a known feedback effect (**Fig. 3A**, right).(38) Hymeglusin treatment also resulted in an increase of RhoA and HMGCR, along with HMGCS1. Similarly, statin treatment caused a significant increase in RhoA, HMGCR, and the emergence of HMGylated FASN. These data suggest that each condition results in a comparable degree of mevalonate flux perturbation. Accordingly, the TMT-based proteomics approach was applied to the cells under the corresponding conditions in four replicates, and the overall quality of the samples was assessed again through immunoblotting (**Fig. 3B,C**). The following mass-spectrometry analysis revealed the relative abundance of 9,881 proteins across all 16 samples without missing values. Principal component analysis showed that replicates of each sample clustered together, indicating consistency among the replicates (**Fig. S2A**). Notably, only the untreated cells showed a distinct separation, while the cells treated with the mevalonate pathway inhibition formed a closely clustered group. For example, proteome profiles from cells treated with Simvastatin and Hymeglusin showed nearly identical results, indicated by a strong Pearson correlation coefficient of 0.97 (**Fig. S2B**). A similarly high correlation coefficient (Pearson r = 0.95) was observed when comparing Hymeglusin-treated cells with those subjected to HMGCS1 degradation (**Fig. S2C**). These unbiased proteomics data reveal that 1) Hymeglusin produces proteome changes that are highly specific to HMGCS1 inhibition, 2) HMGCS1 depletion causes proteome changes driven by the loss of its enzymatic function, and 3) targeting HMGCS1 and HMGCR leads to the nearly identical proteomic changes in 24 hours, reinforcing their essential role in the mevalonate pathway. 12% of the total proteome (1,200 of 9,881 proteins) showed significant changes within 24 hours of Hymeglusin treatment. It was evident that the inhibition of the mevalonate pathway substantially stimulated the production of a subset of the proteome, with 168 proteins increasing by more than two-fold while 24 proteins decreased (**Fig. 3D**). Gene Ontology (GO) analysis of the significantly upregulated proteins (> 2-fold, 168 proteins) for their cellular compartment indicated that the proteins in the secretory pathway are significantly enriched (**Fig. 3E**). Functionally, proteins involved in regulating cell migration and signaling receptor binding were prominent (30 and 38 proteins, respectively), aligning with the role of the mevalonate pathway in regulating small GTPase family members, such as RHO, RAB, and RAS. A notable increase was also observed in ∼ 20 enzymes within the mevalonate pathway across the various treatment conditions (**Fig. 3F**). All three inhibitory conditions resulted in a similar induction of the mevalonate pathway enzymes. These findings are consistent with prior reports of a transcriptional feedback mechanism mediated by Sterol Regulatory Element-Binding Protein 2, which is triggered upon the inhibition of MVP enzymes.(39) Significantly downregulated proteins (greater than 2-fold difference, 24 proteins) included seven proteins involved in the mitotic spindle checkpoint, indicating that cell cycle arrest was initiated, likely as a sign of early-stage apoptosis (**Fig. 3D**). In this context, BCL2L12, an anti-apoptotic regulator, was downregulated by more than 2-fold. Mevalonate is a precursor for not only sterols but also isoprenoids that are essential for post-translational modifications, such as farnesylation and geranylgeranylation. Three proteins targeted by farnesylation (RAB20, RAB29, RAB12) and two proteins for geranylgeranylation with a CAAX motif (MIEN1 and FBXO10) were significantly downregulated, suggesting their destabilization due to the loss of prenylation. When we examined the changes in prenylation substrates on a broader scale, we found many of them among the statistically significant hits. Intriguingly, they showed divergent expression changes upon the inhibition of mevalonate production (**Fig. 3G**). Their downregulation may stem from instability caused by the lack of prenylation in the newly produced proteins, while the up-regulated proteins may be a result of their transcriptional feedback effect mediated by SREBP2, producing unprenylated RHO and RAB proteins. In any case, their cellular functions may be negatively affected by the loss of prenylation. In summary, the inhibition or degradation of HMGCS1, or the inhibition of HMGCR by statins, results in a similar alteration of the global proteome that is characterized by the induction of the MVP enzymes and the substrates of the downstream metabolites, isoprenoids.

**FIGURE 3.**
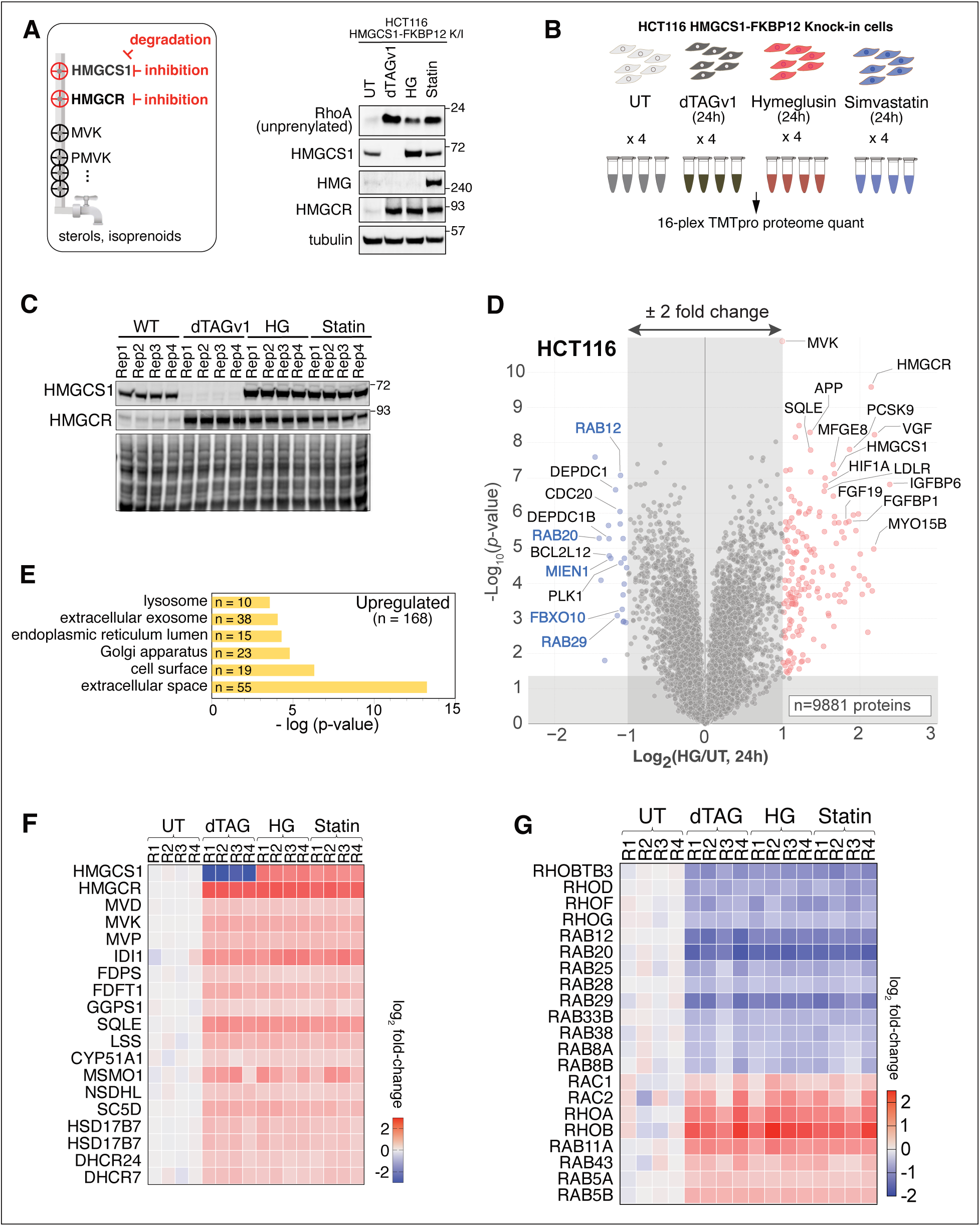
Perturbing HMGCS1 by Hymeglusin and HMGCR by Simvastatin leads to identical global proteome changes. (A) left panel: A schematic illustrating the mevalonate pathway flux. Three perturbation approaches— inhibition or degradation of HMGCS1 and inhibition of HMGCR—are compared for their effects on global proteome changes. Right panel: HCT116 HMGCS1-FKBP12^F36V^ knock-in cells were treated with dTAGv1 (100 nM, for HMGCS1 degradation), Hymeglusin (5 μM, for HMGCS1 inhibition), and Simvastatin (10 µM, for HMGCR inhibition) for 48 hours. The cell extracts were then probed with the indicated antibodies to check the cellular response. (B) The 16-plex TMTpro approach was utilized to compare the global proteome changes after 24 hours of the specified small molecule treatment. (C) HEK293T cells treated as described in panel B were probed with the indicated antibodies as a quality control step prior to proteomic analysis. (D) Analysis of the TMT-plex data is presented as a volcano plot of the —log_10_-transformed p-value versus the log_2_-transformed ratio of Hymeglusin treated/untreated (UT) conditions for HCT116 HMGCS1-FKBP12^F36V^ knock-in cells. p values were calculated by two-sided Welch’s t-test (adjusted to 1% FDR for multiple comparisons). A total of 9881 proteins were quantified. n = 4 biological replicates. (E) Gene ontology analyses of the statistically significant upregulated proteins (168 proteins) with more than 2-fold changes are shown. N represents the number of proteins counted in the category. (F) 19 quantified proteins in the mevalonate/sterol pathway were curated and their log2 fold-changes are presented as a heat map. dTAG treatment led to the steep depletion of HMGCS1. (G) Representative families of isoprenylation targets (Rho, Rab, Rheb, and Ras proteins) were curated from 1,200 statistically significant hits and plotted as a heatmap. No Ras or Rheb proteins showed statistically significant change upon treatment.

### Simultaneous targeting HMGCS1 and HMGCR produces a synergistic anti-cancer response

Repurposing statins has attracted significant interest as a strategy for treating cancer.(20) However, many patients were non-responsive to statin treatment. Prior studies have suggested that cells demonstrating feedback upregulation of the MVP enzymes, particularly HMGCS1, tend to exhibit strong statin resistance.(40–42) We tested this by using seven cell lines with different lineages. Simvastatin treatment for 24 hours led to varying levels of HMGCS1 induction (**Fig. S3A, B**). Among the seven cell lines, HCT116 showed a 2.5-fold increase in HMGCS1, while the HMGCS1 levels in HEK293T or MFE296 cell lines remained unaltered (**Fig. S3C**). The following cell viability assay on HCT116 and HEK293T cells indicated that HCT116 cells are highly resistant to Simvastatin treatment, while HEK293T cells were relatively more sensitive (**Fig. S3D**). Colony formation assays on HCT116 and MFE296 cells also showed resistance for HCT116, whereas MFE296 cells exhibited a more sensitive response (**Fig. S3E**). Based on our data and the prior study, we conclude that the increased production of HMGCS1 upon statin treatment may contribute to the low efficacy of statins in HCT116.

The strong correlation observed between changes in the global proteome caused by the inhibition of HMGCS1 and HMGCR suggests that simultaneously targeting both enzymes may more effectively inhibit the mevalonate metabolic pathway. This could address the current limitation of statin efficacy in anti-cancer therapy, especially in cells that are resistant to statins. We, therefore, examined if simultaneous targeting of HMGCS1 and HMGCR would enhance the anti-proliferative effect of the monotherapy using HCT116 cells (**Fig. 4A**). In a colony formation assay, treatment with a statin alone resulted in a ∼30% decrease in the number of colonies, while Hymeglusin alone had a negligible effect. In contrast, combined treatment with Symvastatin and Hymeglusin resulted in significant inhibition of colony formation. Similarly, we examined the effect of HMGCS1 degradation on cell proliferation, both individually and in combination with a statin, using HCT116 cells expressing endogenous HMGCS1-FKBP12^F36V^ fusion protein (**Fig. 4B**). In these cells, Simvastatin treatment resulted in a substantial increase in endogenous HMGCS1-FKBP12^F36V^, as expected, whereas addition of dTAGv1 nearly depleted HMGCS1 via induced degradation. Treatment of dTAGv1 in these cells led to a 60% reduction in colony numbers, while co-treatment with dTAGv1 and Simvastatin resulted in nearly complete inhibition of colony formation (**Fig. 4C,D)**. Similarly, cell viability assay results gained from the monolayered cell culture system indicated that the addition of dTAGv1 ligand significantly enhanced the IC_50_ of Simvastatin, decreasing from 15.6 µM to 1.3 µM (**Fig. 4E**). As a control, we treated wild-type HCT116 cells with the dTAGv1 ligand, which exhibited no change in the IC_50_ of Simvastatin, confirming that the observed change in IC_50_ was due to the degradation of HMGCS1 (**Fig. 4E**, bottom). Importantly, the cell death caused by the joint treatment was rescued by supplementing the media with geranylgeranyl alcohol (GGOH), a precursor of the downstream metabolite of the mevalonate, indicating that the enhanced cytotoxicity arises from the impairment of the mevalonate synthesis (**Fig. 4F**).

**FIGURE 4.**
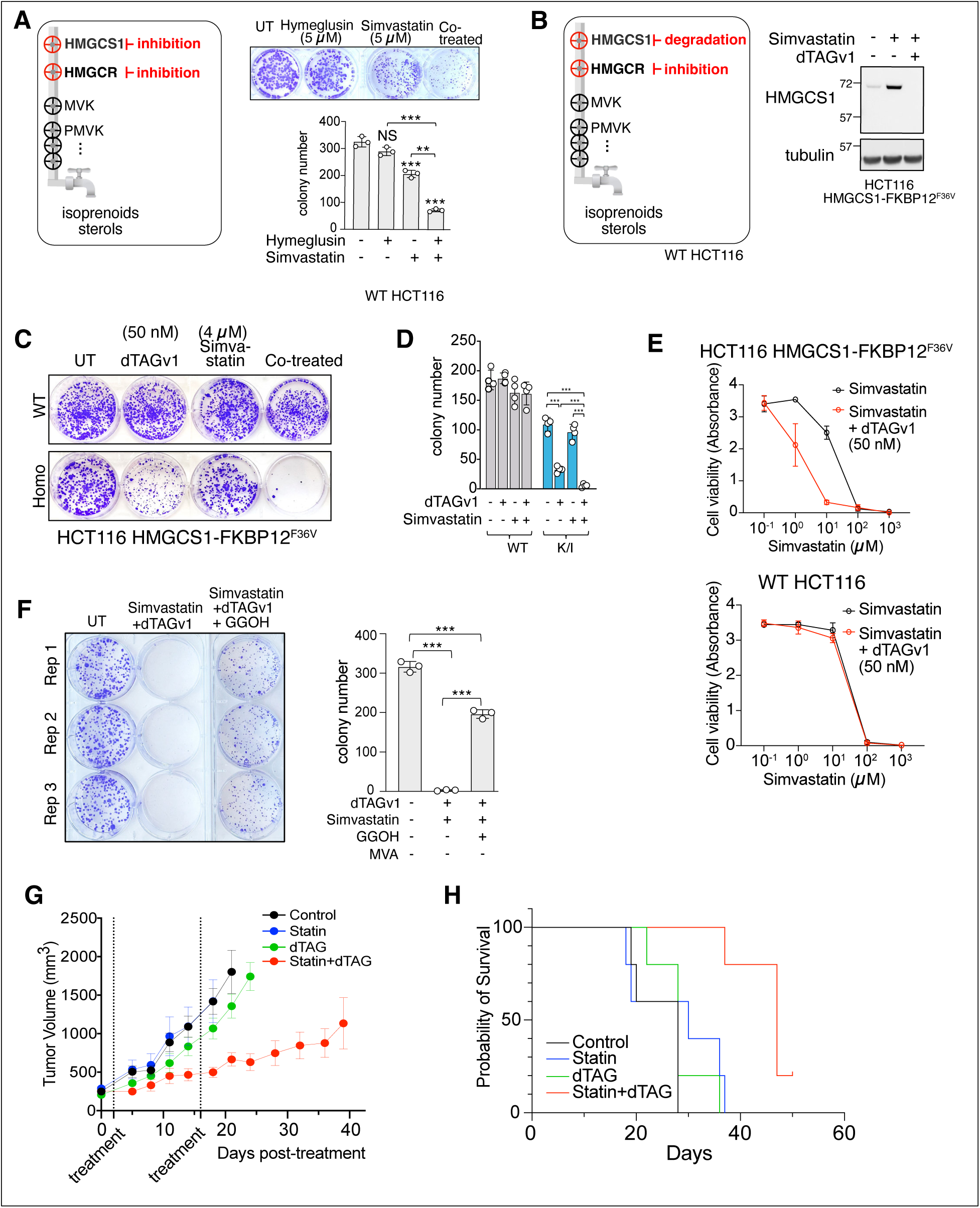
Combinatory targeting HMGCS1 and HMGCR exhibit synergistic anti-tumor effects. (A) The left panel presents a schematic illustration of the combinatorial inhibition of HMGCS1 and HMGCR. In the right panel, the colony formation assay results for HMGCS1-FKBP12^F36V^ K/I HCT116 cells are shown, where treatments involved either Hymeglusin (5 μM), Simvastatin (5 μM), or a combination of both inhibitors. Corresponding inhibitors were applied on days 5 and 7 after seeding, and the colony numbers were counted on day 12. The accompanying quantification graph displays the means ± SD from biological triplicates. (B) The left panel illustrates the degradation of HMGCS1 and the inhibition of HMGCR as a combinatorial approach to target the mevalonate pathway. The right panel shows the induction of HMGCS1 following Simvastatin treatment, which was completely reversed by dTAGv1 treatment due to the induced degradation in HCT116 cells expressing endogenous HMGCS1-FKBP12^F36V^. (C) HMGCS1 degradation by dTAGv1 (50 nM) increases the inhibitory effects of Simvastatin (4 μM) on the colony-forming activity of HCT116 HMGCS1-FKBP12^F36V^ cells. (D) Means ± s.d. of biological quadruplicate data from Panel C. (E) HMGCS1 degradation by the dTAG system reduces the IC_50_ value of Simvastatin by 10-fold in HCT116 HMGCS1-FKBP12^F36V^ knock-in cells (top). Cell viability was assessed 24 hours after treating the corresponding cells with increasing concentrations of Simvastatin and dTAGv1 (50 nM). The wild-type HCT116 did not show the synergistic effect of dTAGv1 and Simvastatin (bottom). (F) The anti-colony formation effect induced by the HMGCS1 degradation and HMGCR inhibition was rescued by supplementation of geranylgeraniol (GGOH, 40 μM). HCT116 HMGCS1-FKBP12^F36V^ K/I cells were treated with dTAGv1 (50 nM) and Simvastatin (4 μM) on days 5 and 7, post-seeding. GGOH (40 µM) was added to the media of the corresponding cells on days 5 and 7. Means ± s.d. of biological triplicate data is shown on the right. (G) HMGCS1-FKBP12^F36V^ K/I HCT116 cells were implanted into nude mice. When the tumor size reached 200 – 250 mm^3^, we administrated two treatment regimens (shown in dotted vertical lines), including saline control, 5 mg/kg Simvastatin via gavage three times a week, weekly 5mg/kg dTAGv1 through i.p., and the combination of both dTAGv1 and Simvastatin. Degradation of HMGCS1 by the dTAG system potentiates the tumor growth-suppressing effect of Simvastatin. n = 5 mice for each condition. (H) Survival probability analysis of the mice presented in Panel H using Kaplan-Meier software reveals extended mortality in the dTAG and Statin co-treatment groups compared to the other control groups.

Lastly, we performed the equivalent study using the mouse xenograft model after implanting HCT116 HMGCS1-FKBP12^F36V^ cells (**Fig. 4G,H**). Four groups of mice were subjected to two treatment regimens over a two-week interval, including saline control, oral Simvastatin three times a week, weekly i.p. injection of dTAGv1, and the combination of Simvastatin and dTAGv1. The group of mice that received both Simvastatin and dTAGv1 exhibited a synergistic effect on tumor growth suppression that lasted for 40 days. In contrast, administration of Simvastatin or dTAGv1 ligand injection alone did not show a significant difference compared to the PBS-treated control group. Overall data suggest that dual targeting of HMGCS1 and HMGCR may offer potential benefits, and the targeted proteolysis approach for HMGCS1 perturbation could demonstrate greater efficacy compared to inhibition.

### Hymeglusin warhead exhibits poor serum stability

Despite identical proteome changes upon degradation or inhibition of HMGCS1 (**Fig. 3**), enzymatic inhibition of HMGCS1 by Hymeglusin did not elicit any cytotoxic effects in either HCT116 or HEK293T cells. when used alone (**Figs. 4A and S4A**). This observation is inconsistent with the anti-tumor efficacy of dTAGv1-mediated HMGCS1 degradation, prompting us to further investigate the long-term stability of Hymeglusin. Prior investigations using murine models administered with Hymeglusin reported an 80% reduction in cholesterol synthesis within 30 min and a complete loss of inhibitory function by 180 min when measured by the conversion of ^14^C-labeled acetate into plasma cholesterol.(25, 26) It was speculated that this effect may be due to the reversible nature of the thioester bond formed between Hymeglusin and HMGCS1. However, the precise kinetics underlying the decline in Hymeglusin efficacy and the mechanism behind its loss of efficacy in cells remain to be determined. Considering that researchers have utilized Hymeglusin as an inhibitor of HMGCS1 in a range of cell culture models, elucidating this limitation could be crucial for its appropriate application. Our use of the HMGCS1 activity-based probes enabled us to monitor the occupancy of the catalytic cysteine in HMGCS1 by Hymeglusin over time. After 8 hours of treatment with 0.5 µM Hymeglusin, we observed the emergence of the catalytically active HMGCS1 that was labeled with HG-FL, a trend that became more pronounced after 16 hours in HEK293T, HCT116, or HepG2 cells (**Fig. 5A,B**). These data indicate that HMGCS1 labeling is not maintained over the long term, which aligns with the absence of sustained efficacy in the antiproliferative effects we observed with Hymeglusin treatment. Interestingly, the removal of serum from the culture media led to a delayed restoration of active HMGCS1 levels, indicating that serum plays a role in the short-term efficacy of Hymeglusin (**Fig. 5C**). Indeed, when Hymeglusin was pre-incubated in DMEM with 10% FBS for 16 hours before being added to the cells, the catalytic cysteine of HMGCS1 was not effectively inhibited anymore (**Fig. 5D,E**). In contrast, pre-incubation in DMEM without 10% FBS did not reduce the efficacy of Hymeglusin. Additionally, we observed intense labeling of bovine serum albumin (BSA) by HG-

**FIGURE 5.**
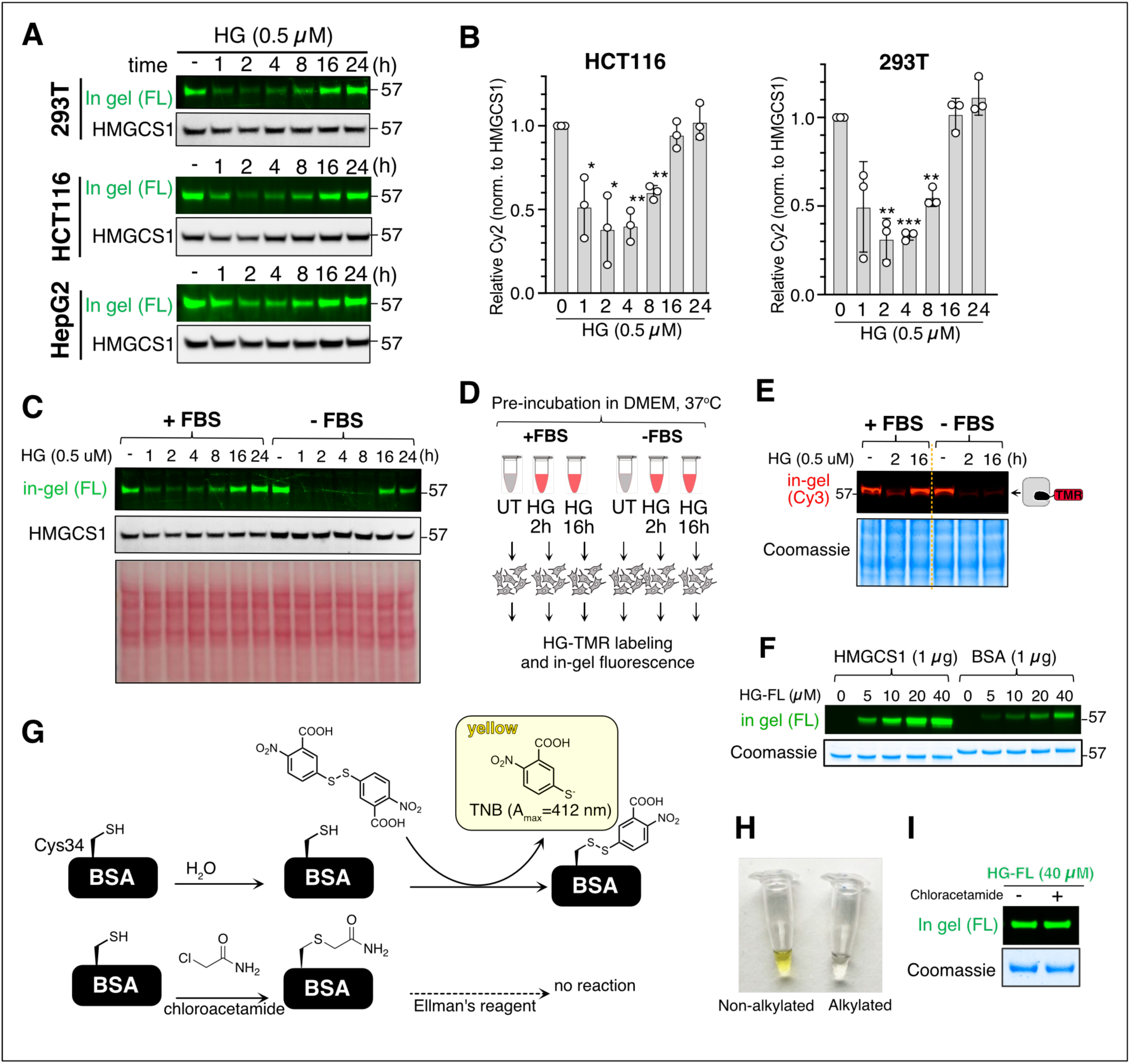
Hymeglusin exhibits poor serum stability. (A) HCT116, HEK293T, and HepG2 cells were treated with Hymeglusin (0.5 µM) for the indicated time points, followed by lysis and reaction with the HG-FL probe. In-gel fluorescence analysis and immunoblotting were then performed (B) The quantification of relative fluorescence intensity from three replicate experiments in panel A is presented. Mean ± s.d. (C) The absence of FBS in the cell culture media delays the appearance of active HMGCS1 with free catalytic cysteine after prolonged treatment of cells with Hymeglusin. (D) A workflow to test the effect of fetal bovine serum (FBS) on the stability of Hymeglusin. 0.5 μM of Hymeglusin was added to DMEM supplemented with or without 10% FBS. The media was then incubated at 37°C for 2 or 16 hours before being added to HEK293T cells. (E) HEK293T cells treated as in panel D were subjected to in-gel fluorescence analysis. (F) *In vitro* incubation of HG-FL with recombinant HMGCS1 (1 µg) or BSA (1 µg) for 1 hour produces a green fluorescence signal in the relevant molecular weight regions, indicating a strong interaction between HG-FL and BSA. (G) Scheme of Ellman’s reagent, which turns the solution into a yellow visible color when reacted with thiol by producing TNB. (H) A portion of the samples treated as in G were reacted with Ellman’s reagent. Specifically, BSA was incubated with water or chloroacetamide for 10 minutes, followed by the addition of Ellman’s reagent. The water pre-treated sample turned into the expected yellow solution, while the chloroacetamide pre-treated BSA solution did not change color, indicating the loss of free thiol due to alkylation. (I) The remaining portion of the samples in panels G and H was then incubated with HG-FL, which showed an equal level of green fluorescence signal.

FL after incubating them for one hour *in vitro* (**Fig. 5F**). The labeling persisted even after boiling for 5 minutes in denaturing conditions, suggesting a strong interaction between HG-FL and BSA. We tried to clarify this by categorizing the potential pathways that underlie how BSA impacts the efficacy of Hymeglusin into two distinct types: First, BSA contains 35 cysteines, of which 34 form disulfide bonds, leaving one reactive cysteine.(43) Thus, Hymeglusin may form a covalent bond with BSA via reactive Cys34. The second possibility is that Hymeglusin forms a non-covalent, yet strong, interaction through the fatty acid binding sites, as BSA has several fatty acid binding sites and Hymeglusin resembles the structure of fatty acids.(44) As the second possibility is rather complex to address, we evaluated the first hypothesis. If the cysteine on the BSA is reacting with HG-FL, then alkylating BSA with chloroacetamide prior to HG-FL incubation should result in no labeling. We thus treated BSA with chloroacetamide, and the loss of reactive cysteine was confirmed by using Ellman’s reagent (**Fig. 5G, H**). When native BSA was incubated with Ellman’s reagent, the solution changed to yellow due to the formation of 2-nitro-5-thiobenzoate (TNB). After pre-incubating the BSA with chloroacetamide, we observed the expected loss of color change, representing the absence of active cysteine on BSA (**Fig. 5H**). Then, this alkylated BSA was treated with HG-FL, with non-alkylated BSA serving as a positive control (**Fig. 5I**). The subsequent in-gel fluorescence analysis revealed an unchanged fluorescence signal, suggesting that hymeglusin likely binds to BSA through non-covalent interactions. Together, poor serum stability may limit the long-term use of Hymeglusin in cell and in vivo studies, as human serum typically contains ∼ 40 mg/mL of albumin.

## DISCUSSION

The mevalonate pathway is a well-established focus of interest for chemists and biologists, given its critical role in cholesterol biogenesis and prenylation. Prenylation, in particular, is essential for the proper function of several oncogenic proteins, such as RAS, which underscores its contribution to cancer progression. Consequently, several enzymes in the mevalonate pathway have received substantial attention in pharmaceutical research for inhibitor development. However, the initial enzyme of the pathway, HMGCS1, has received relatively little focus, overshadowed by HMGCR and the established use of statins. We have recently discovered that HMGCS1 functions as a gatekeeper of this pathway, especially in cells with hyperactive mTORC1 signaling.(22) HMGCS1 senses and responds to fluctuations in mTORC1 activity, paralleling the critical role of HMGCR in sensing and responding to changes in membrane sterol levels.

Moreover, the presence of a ligandable cysteine in HMGCS1 makes it a particularly unique and advantageous target for probe development, which can significantly facilitate investigations into both the mechanisms of inhibitor action and potential resistance.

Currently, few tools are available to interrogate HMGCS1. Among them, Hymeglusin, a natural product derived from fungi, stands out as the primary tool utilized in the field since its discovery in the 1980s.(25–29, 45) Despite its widespread use, the selectivity and overall efficacy of Hymeglusin in cells have not been thoroughly evaluated since the 1990s, raising concerns about its suitability as a tool compound. This prompted us to assess its cellular potency and selectivity by developing activity-based probes inspired by the Hymeglusin warhead and leveraging modern technologies, such as proteomics. Our unbiased analyses, grounded in chemical biology approaches, revealed that Hymeglusin is a potent and highly selective inhibitor of HMGCS1 in HEK293T cells, with a concentration of 0.5 µM fully saturating the catalytic cysteine of HMGCS1 in cells. However, we also determined that Hymeglusin has extremely poor serum stability, apparently derived from the interaction with serum albumin, which is comparable to on-target binding to HMGCS1. This drawback limits the duration and efficacy of Hymeglusin to target HMGCS1. Nevertheless, our comprehensive data led us to consider Hymeglusin a viable tool for studying the short-term pharmacologic effects of HMGCS1 inhibition, provided we carefully monitor its intracellular stability and activity with our activity-based probes.

After carefully characterizing Hymeglusin, we explored the effect of inhibiting HMGCS1 using Hymeglusin and an additional method that incorporates a chemical genomic approach to induce the degradation of HMGCS1. Proteomic analysis of cells following HMGCS1 inhibition or degradation, or HMGCR inhibition, revealed surprisingly similar proteome changes in 24 hours. This result indicates that targeting both enzymes simultaneously may enhance drug efficacy without amplifying the risk of potential side effects from either inhibitor. Our follow-up cell biological and xenograft studies support this notion that dual targeting of HMGCS1 and HMGCR potentiates the anti-cancer effect of statin monotherapy. Overall, our findings highlight the potential of targeting HMGCS1 either as a monotherapy or in combination with statins, emphasizing the benefits of the induced proteolysis approach in this context.

Comparative analysis of HMGCS1 inhibition by Hymeglusin or of HMGCS1 degradation using the chemically induced tag system (dTAG) demonstrated the superior potency of the degradation approach in inhibiting cancer cell proliferation. Heterobifunctional degrader molecules have potential therapeutic advantages over the monofunctional inhibitor.(46, 47) In the case of HMGCS1, the proteolysis targeting chimera (PROTAC) degrader can counteract the compensatory upregulation of HMGCS1 through its degradation. As the half-life of the HMGCS1 protein spans several days, the drug-induced degradation of HMGCS1 can be prolonged longer than the inhibitors. Additionally, covalent PROTACs have been shown to enhance cellular uptake and target selectivity.(48) Therefore, developing small molecules that could induce targeted proteolysis of HMGCS1 may offer better pharmacological properties than inhibitors, providing a more effective strategy to treat cancer both as monotherapy and in combination with HMGCR inhibitors.

Although we here defined that Hymeglusin interacts with albumin through non-covalent binding, further investigation is needed to determine the detailed binding mode between Hymeglusin and albumin protein.(49) One possibility is that the structure of Hymeglusin resembles that of fatty acids. Albumin has well-characterized fatty acid binding pockets, with at least seven known pockets that could potentially interact with Hymeglusin.(50) Another limitation of the present study is that the mitochondrial HMG-CoA synthase, HMGCS2, which shares approximately 60% sequence identity with HMGCS1, may be another potential target of Hymeglusin. HMGCS2 exhibits tissue-specific expression profiles, unlike the universal expression of HMGCS1. Therefore, this aspect should be considered when using Hymeglusin for cell lines that express HMGCS2.

In summary, this study reports new chemical biology tools for an important yet underexplored metabolic enzyme within cells, HMGCS1. It offers insights into how targeting HMGCS1—either alone or together with HMGCR—can lead to alterations in the identical cellular proteome while potentiating their anti-proliferation effect. Furthermore, the study holds potential for developing drug-like inhibitors or bisubstrate degraders of HMGCS1 by utilizing the activity-based probes developed in this study.

## EXPERIMENTAL PROCEDURES

Detailed information about reagents and resources used in this study are displayed in Table S3.

### Cell culture

HEK293T, HCT116, RPE1, HeLa, and HepG2 cells were grown in Dulbecco’s modified Eagle’s medium (DMEM), supplemented with 10% fetal bovine serum and maintained in a 5% CO_2_ incubator at 37°C.

### Cell lysis and immunoblotting assay

Cells were plated 24 h before being treated with inhibitors or starvation medium, and cell confluency did not exceed 70% at the time of harvest. Cell pellets were lysed with either in-house RIPA buffer (50 mM HEPES, 150 mM NaCl, pH 7.6, 1% NP-40, 1% sodium deoxycholate, 0.1% SDS, 10 mM glycerophosphate, 10 mM sodium pyrophosphate, protease inhibitor cocktail, phosphatase inhibitor cocktail) containing 200 μM TCEP, 15 mM MgCl_2,_ and benzonase (Millipore, for RIPA only) or 0.5 % NP40 buffer (50 mM HEPES HCl, 150 mM NaCl, pH 7.4) containing protease inhibitors, phosphatase inhibitors, and 200 μM TCEP. Lysates in RIPA buffer were then sonicated, whereas lysates in NP40 buffer were pipetted and centrifuged at 7,000 rpm for 1 min without sonication. Bradford assay was performed to measure the protein concentration, and the normalized cell lysates were denatured by adding LDS supplemented with 50 mM DTT, followed by boiling at 90 °C for 5 minutes. Less than 30 μg of each lysate was loaded onto the 4-12% NuPAGE Bis-Tris gel (Thermo Fisher Scientific), followed by SDS-PAGE with MES running buffer. Following transfer to PVDF membranes (0.45 μm or 0.2 μm, Millipore), blocking was performed with 5% non-fat milk at room temperature for 15 min, then incubated with primary antibodies overnight at 4°C. After the incubation with primary antibobies, membranes were washed with TBST and further incubated with fluorescent IRDye antibody (1:20000) for 1 h at room temperature. The membrane was then imaged using Chemidoc MP (Biorad).

### Total proteomics analysis using TMTpro

The total proteomics analyses were performed based on the previously reported method.(22) Briefly, cells are plated into a 10 cm dish per condition, a total of 16 dishes. 24 h later, dTAGv1 (100 nM), Hymeglusin (5 µM), or Simvastatin (10 µM) were added to cells and incubated for 24 1. h. Cells were then washed with ice-cold PBS five times and lysed with 800 μl of RIPA buffer (50 mM HEPES, 150 mM NaCl, pH 7.6, 1% NP-40, 1% sodium deoxycholate, 0.1% SDS, 10 mM glycerophosphate, 10 mM sodium pyrophosphate, protease inhibitor cocktail, Phosphatase inhibitor cocktail, pH 7.5). The lysate was then collected and sonicated three times, followed by Bradford assay to measure the protein concentration. The total protein concentration was adjusted to become 3 mg/ml throughout the samples using RIPA buffer. 125 μg of each protein extract was taken and reduced by incubation in the presence of 5 mM TCEP at 55 °C for 10 min. After cooling down to room temperature, the lysates were reacted with chloroacetamide solution (final conc. 20 mM) at room temperature for 15 min, followed by chloroform/methanol precipitation. Protein discs were resuspended in 100 mM EPPS (pH 8.5) containing 0.1% RapiGest and digested at 37 °C overnight with Trypsin (100:1 protein-to-protease ratio). 16-plex tandem mass tag labeling of each sample was performed by adding 10 µL of the 20 ng/mL stock of TMTpro reagent along with acetonitrile to achieve a final acetonitrile concentration of approximately 30% (v/v). Following incubation at room temperature for 1 h, the reaction was quenched with hydroxylamine to a final concentration of 1% (v/v) for 15 min. The TMTpro-labeled samples were pooled together at a 1:1 ratio. The sample was vacuum centrifuged to near dryness and subjected to C18 solid-phase extraction (SPE) (50 mg, Sep-Pak, Waters). Dried TMTpro-labeled sample was resuspended in 100 μL of 10 mM NH_4_HCO_3_ pH 8.0 and fractionated using BPRP HPLC. Briefly, samples were offline fractionated over a 90 min run into 96 fractions by high pH reverse-phase HPLC (Agilent LC1260) through an aeris peptide xb-c18 column (Phenomenex; 250 mm x 3.6 mm) with mobile phase A containing 5% acetonitrile and 10 mM NH_4_HCO_3_ in LC-MS grade H_2_O, and mobile phase B containing 90% acetonitrile and 10 mM NH_4_HCO3 in LC-MS grade H_2_O (both pH 8.0). The 96 resulting fractions were then pooled in a non-continuous manner into 24 fractions (as outlined in Figure S5 of Paulo et al., 2016)(51) for mass spectrometry analysis. Fractions were vacuum centrifuged to near dryness. Each consolidated fraction was desalted via Stage Tip, dried again via vacuum centrifugation, and reconstituted in 5 % acetonitrile, 1 % formic acid for LC-MS/MS processing.

Mass spectrometry data were acquired using an Orbitrap Eclipse Tribrid mass spectrometer (Thermo Fisher Scientific, San Jose, CA) connected to an UltiMate 3000 RSLCnano system liquid chromatography (LC) pump (Thermo Fisher Scientific, San Jose, CA). Peptides were separated on a 100 μm inner diameter microcapillary column packed in-house with ∼ 30 cm of HALO Peptide ES-C18 resin (2.7 μm, 160 Å, Advanced Materials Technology, Wilmington, DE) with a gradient consisting of 5 %–23 % (0-75 min), 23-40 % (75-110 min) (ACN, 0.1% FA) over a 120 min run at ∼500 nL/min. 3/10 of each fraction was loaded onto the column for analysis. Proteome analysis used Multi-Notch MS^3^-based TMT quantification, combined with Real Time Search analysis software, and the FAIMS Pro Interface (using previously optimized 3 CV parameters), to reduce ion interference. The scan sequence began with an MS^1^ spectrum (Orbitrap analysis; resolution 120,000 at 200 Th; mass range 400−1500 m/z; maximum injection time 50 ms; automatic gain control (AGC) target 4×10^5^). For MS2 analysis, precursors were selected based on a cycle time of 1.25 sec/CV method (FAIMS CV=-40/-60/-80). MS^2^ analysis consisted of collision-induced dissociation (quadrupole ion trap analysis; Rapid scan rate; AGC 1.0×10^4^; isolation window 0.5 Th; normalized collision energy (NCE) 35; maximum injection time 35 ms). Monoisotopic peak assignment was used, and previously interrogated precursors were excluded using a dynamic window (180 s ±10 ppm). Following the acquisition of each MS^2^ spectrum, a synchronous-precursor-selection (SPS) API-MS^3^ scan was collected on the top 10 most intense ions b or y-ions matched by the online search algorithm in the associated MS^2^ spectrum. MS^3^ precursors were fragmented by high energy collision-induced dissociation (HCD) and analyzed using the Orbitrap (NCE 45; AGC 2.5×10^5^; maximum injection time 200 ms, resolution was 50,000 at 200 Th). Closeout was set at two peptides per protein per fraction, so that MS^3^s were no longer collected for proteins having two peptide spectrum matches (PSMs) that passed quality filters.

### Proteomics Data Analysis

Mass spectra were processed using a Comet-based (2020.01 rev. 4) software pipeline (52, 53). Spectra were first converted to mzXML and monoisotopic peaks were re-assigned using Monocle software (54). MS^2^ spectra were matched with peptide sequences with a composite sequence database including the Human Reference Proteome (2020-01 - SwissProt entries only) UniProt database, as well as sequences of common contaminants. This database was concatenated with one composed of all protein sequences in the reversed order. Analysis was performed using a 50 ppm precursor ion tolerance. Static modifications included TMTpro tags on lysine residues and peptide N termini (+304.207 Da) and carbamidomethylation of cysteine residues (+57.021 Da). Oxidation of methionine residues (+15.995 Da) was set as a variable modification. Peptide-spectrum matches (PSMs) were adjusted to a 1% false discovery rate (FDR) (55). PSM filtering was performed using a linear discriminant analysis, (56), while considering the following parameters: Comet log expect, different sequence delta Comet log expect, missed cleavages, peptide length, charge state, precursor mass accuracy, and fraction of ions matched. For protein-level comparisons, PSMs were identified, quantified, and collapsed to a 1% peptide false discovery rate (FDR) and then collapsed further to a final protein-level FDR of 1% (57). To generate the smallest set of proteins required to account for all observed peptides, the principles of parsimony were applied. For TMTpro-based reporter ion quantitation, the summed signal-to-noise (S:N) ratio for each TMT channel was first extracted based on the closest matching centroid to the expected mass of the TMT reporter ion (integration tolerance of 0.003 Da). Isotopic impurities of the different TMTpro reagents provided by the manufacturer specifications, were used to adjust reporter ion intensities. Proteins were quantified by summing reporter ion signal-to-noise measurements across all matching PSMs, resulting in a ‘‘summed signal-to-noise’’ measurement. For total proteome, PSMs with poor quality, or isolation specificity less than 0.75, or with TMT reporter summed signal-to-noise ratio that were less than 160 or had no MS^3^ spectra were excluded from quantification. For phospho proteome, PSMs with poor quality, MS^2^ spectra with 13 or more TMT reporter ion channels missing, or isolation specificity less than 0.8, or with TMT reporter summed signal-to-noise ratio that were less than 160 were excluded from quantification. The AScore algorithm was used to determine the localization of phosphorylation sites (58). AScore is a probability-based approach for high-throughput protein phosphorylation site localization. Precisely, a threshold of 13 relates to 95% confidence in site localization.

Protein quantification values were exported for further analysis in Microsoft Excel and Perseus (59). Each reporter ion channel was summed across all quantified proteins and normalized, assuming equal protein loading was achieved for all samples. The maximum and minimum TMT ratio quantifiable was capped at 100-fold. Organellar protein marker annotations were compiled using the proteins that had scored with confidence ‘‘very high’’ or ‘‘high’’ from a previously published HeLa dataset (60) and additional entries from manually curated literature.

Supplemental data tables list all quantified proteins and the TMTpro reporter ratio associated with control channels used for quantitative analysis.

### IP-MS analysis using Hymeglusin-biotin probe

HEK293T cells were seeded overnight before incubating with media containing 5 µM hymeglusin or vehicle control (DMSO) for 0.5 hrs. Then 5 µM Hymeglusin-biotin was introduced to the cells and incubated for additional 2 hrs. Cells were washed with ice-cold PBS three times. The cells were then lysed with 200 μl of HEPES-NP40 buffer (50 mM HEPES, 150 mM NaCl, pH 7.5, 0.25%NP40). The lysate was centrifuged at 7k for 1 min, and the supernatant was collected. The supernatant was then reduced by 5 mM TCEP (10 min, room temperature) and alkylated by chloroacetamide (20 mM final concentration, 15 min, room temperature). MeOH/CHCl_3_ precipitation was followed to remove any remaining Hymeglusin-biotin in the cell lysates. The white protein disk was then resuspended in 2% SDS (50 mM HEPES 150 mM NaCl, pH7.2, 2.5 mM TCEP) and sonicated once with a tip sonicator and then diluted in HEPES-NP40 buffer (final SDS concentration is < 0.5%). Subsequently, 10 µl of high-capacity streptavidin was added to the cell lysates and incubated for 2 hours at r.t.. Flow-through was stored for the quality control immunoblots, and the beads were washed with RIPA X 2, 2%SDS X 3, RIPA X 2, and PBS X 3. The beads were transferred onto hydrophilic PTFE membrane filter cups and washed twice with water. Completely dried beads were resuspended in 50 μl of hexafluoroisopropanol (HFIP), incubated for 5 min with shaking, and eluted by spinning at 5k rpm. 50 μl of HFIP was added to the beads again, and the eluates were combined and dried in speed vac. 6-plex TMT labeling was performed as described in the proteomics sample preparation section.

### In-gel fluorescent analysis with HG-fluorescence probes

Recombinant HMGCS1 protein (1 ug) or cell lysates were incubated with 5 µM Hymeglusin-FITC (HG-FL) or Hymeglusin-TMR (HG-TMR) probe at 37°C for 1 h. The labeling reaction was stopped by adding 4X LDS containing DTT and boiling at 90°C for 5 m. The samples were subjected to SDS-PAGE, and the fluorescent signal in the gel was detected by Chemidoc MP (Biorad).

### Cell viability assay

Cells viability was evaluated using WST1-based colorimetric assay (Roche). 2 × 10^4^ cells in 100 μl of culture medium were seeded for each well of 96 well plates. Next day, the cells were treated with culture media containing various amounts of simvastatin and 50 nM of dTAGv1. After 24 h, cells were incubated with WST-1 solution, which was diluted to 1:10 with culture medium in the cell culture incubator. After 4 h, the absorbance of samples was measured using a microplate reader. The wavelength to measure the absorbance of formazan product from viable cells was 450 nm, and the reference wavelength was 690 nm.

### Colony formation assay

To evaluate the ability of single cancer cells to form a colony, cells were plated at a concentration of 1,500 cells/well onto 6 well plates (day 0). At day 5 and day 7, the cells were treated with the corresponding chemicals. On day 12, the cells were stained with a crystal violet working solution, which contains 0.5% crystal violet and 4% paraformaldehyde in PBS. Numbers of colonies were counted for each well (n = 2 or 3) and presented as mean ± s.d.

### Animal Experiments

Animal experiments were performed in compliance with protocols approved by the Research Animal Resource Center and Institutional Animal Care and Use Committee at Memorial Sloan Kettering Cancer Center. 20 Athymic nude mice (nu/nu athymic mice, Charles River Laboratory) were implanted with HCT116 HMGCS1-FKBP12^F36V^ homozygous knock-in (KI) cells through subcutaneous injection of 2 × 10^6^ cells per mouse. The treatment was initiated as the tumor size became 200-250 mm^3^. Four groups of mice were administered with two treatment regimens over a two-week interval, including saline control, 5mg/kg Simvastatin three times per week via oral gavage, 5 mg/kg dTAGv1 weekly via intraperitoneally injected (IP), and the combination of both Simvastatin and dTAGv1. PBS is used as a vehicle for both simvastatin and dTAGv1. Tumor growth was monitored and recorded twice a week for 40 days, and five mice were used per condition.

### Quantification and statistical analysis

P values in the volcano plots of the proteomics data were analyzed by two-sided Welch’s t-test which was adjusted for multiple comparisons. Comparisons of the quantified data (immunoblotting, colony formation, tumor spheroids) were performed by unpaired Student’s t-test; Statistical significance was judged based on p-values; *p < 0.05; **p < 0.01; ***p < 0.001.

## DATA AND CODE AVAILABILITY

- All unique identifiers, and web links for publicly available datasets are presented within the main text or the supplementary materials. The proteomic data generated in this paper will be deposited to the MassIVE repository with the dataset identifier: MSV000098111
- This paper does not report any original code.
- Any additional information required to reanalyze the data reported in this paper is available from the lead contact upon request.

## ACKNOWLEDGMENTS

We thank Brittany Pham for helping with the cell viability assay. We thank Beth Moorefield, Ph. D., for providing critical feedback and review of the manuscript. We thank Mr. William H. Goodwin and Mrs. Alice Goodwin and the Commonwealth Foundation, and the Center for Experimental Therapeutics at Memorial Sloan Kettering Cancer Center for their support.

## AUTHOR CONTRIBUTIONS

Conceptualization, S.A.Y., L.S. and H.A.; methodology, S.A.Y., L.S., Y.R., and A.O.; Investigation, S.A.Y., L.S., Y.R., and A.O.; writing—original draft, S.A.Y., L.S., and H.A.; writing—review & editing, Y.R., A.O. and J.L.; funding acquisition, A.O., J.L., and H.A.; supervision, H.A.

## FUNDING AND ADDITIONAL INFORMATION

This work was supported by the MSKCC Josie Robertson Foundation (H.A.), NIH R35 CA232130 (J.S. L), and partly by the Memorial Sloan Kettering Cancer Center Support Grant P30CA008748 (A.O., J.S.L., and H.A.)

---

**SI FIGURE 1.**
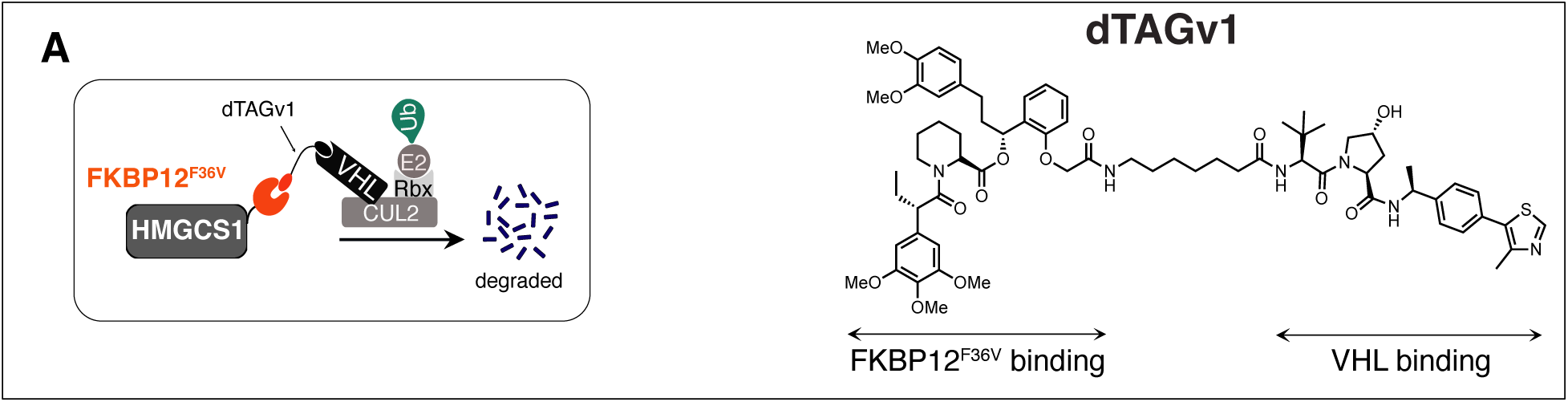
Overview of induced proteolysis by the dTAG system (Related to Figure 1). (A) The schematic of the dTAG system (left) and chemical structure of the dTAGv1 ligand (right). This bifunctional ligand can bring VHL closer to the FKBP12^F36V^ mutant.

**SI FIGURE 2.**
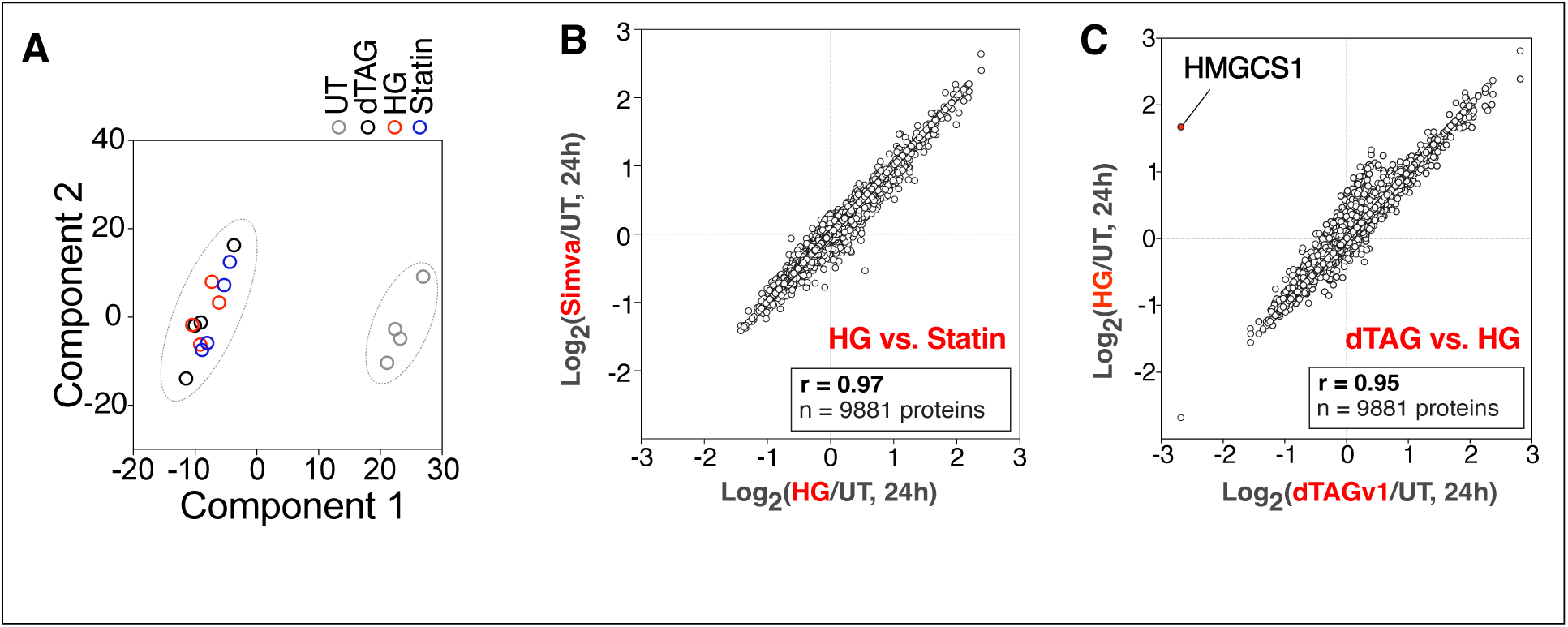
Global proteomic changes induced upon MVP inhibition (Related to Figure 3). (A) Principal component analysis (PCA) of the total proteomics dataset, prepared as shown in Fig. 3B, indicates that dTAG, HG, and Statin-treated cells are grouped together. (B,C) 9881 proteins were plotted for the log_2_-ratio of (Simvastatin/DMSO) against the log_2_-ratio of (HG/DMSO) for panel B, and log_2_-ratio of (Hymeglusin/DMSO) against the log_2_-ratio of (dTAGv1/DMSO) for panel C. N = 4 replicates. p values were calculated by two-sided Welch’s t-test (adjusted to 1% FDR for multiple comparisons). A total of 9881 proteins were quantified. n = 4 biological replicates.

**SI FIGURE 3.**
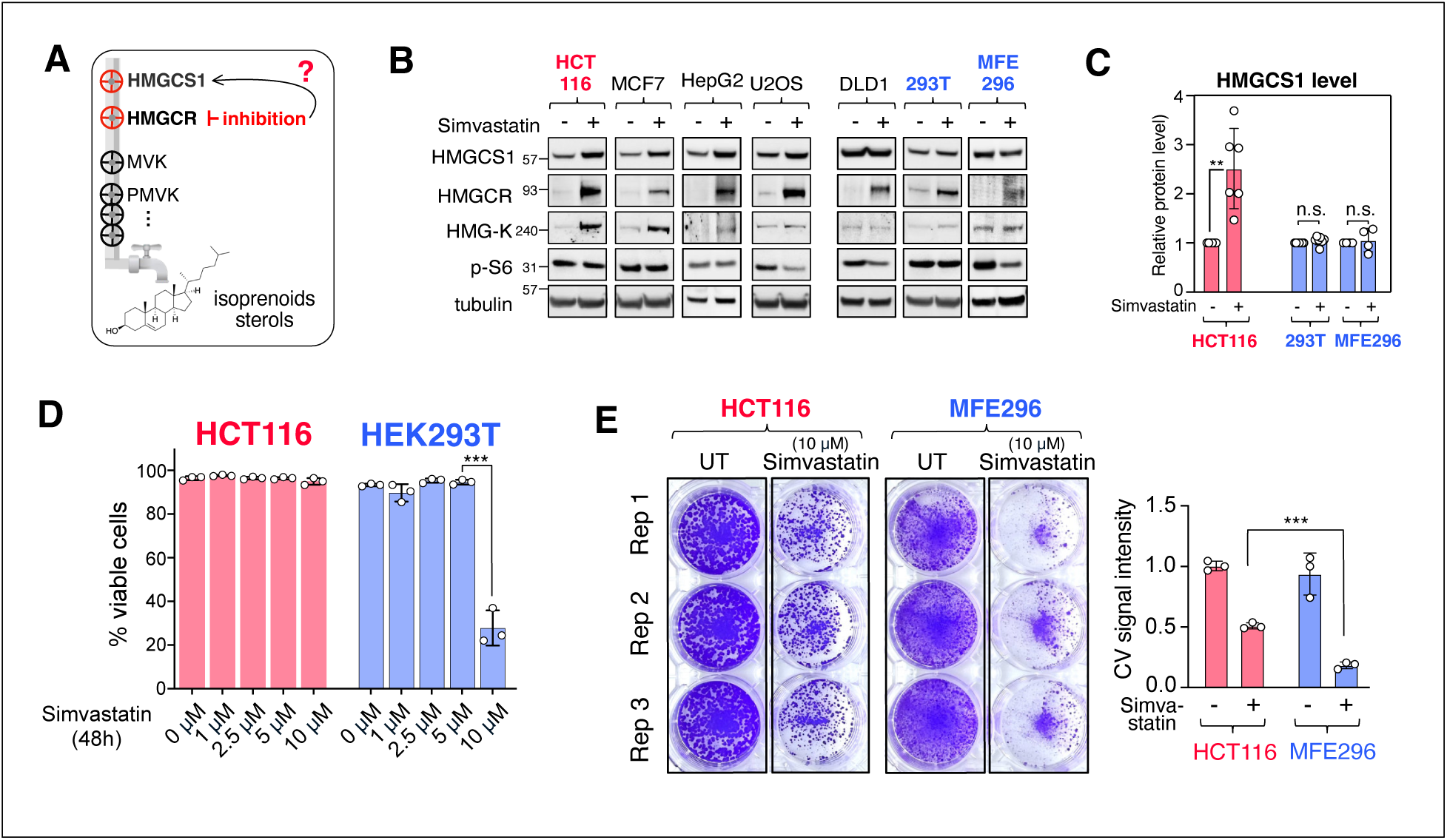
Relation between HMGCS1 upregulation and statin resistance (Related to Figure 4). (A) A schematic showing the relationship between HMGCR inhibition and HMGCS1 up-regulation. (B) The level of HMGCS1 upon Simvastatin treatment for 24 hours was assessed in seven different cell lines using immunoblotting analysis. Indicated cells were treated with Simvastatin (10 μM, 24h), followed by immunoblotting with indicated antibodies. (C) The relative level of HMGCS1 protein was quantified from the experimental replicates shown in panel B, with n > 4 replicates. Mean ± s.d. (D) Flow cytometry analysis of DAPI-stained versus unstained cells was conducted to evaluate cell viability after treating indicated cells with increasing concentrations of Simvastatin for 48 hours. (E) Coloniformation assay with or without Simvastatin treatment was performed on MFE296 and HCT116 cell lines. The total signal intensity per plate, relative to the untreated condition, is presented on the right. Mean ± s.d. n = 3 replicates.

**SI FIGURE 4.**
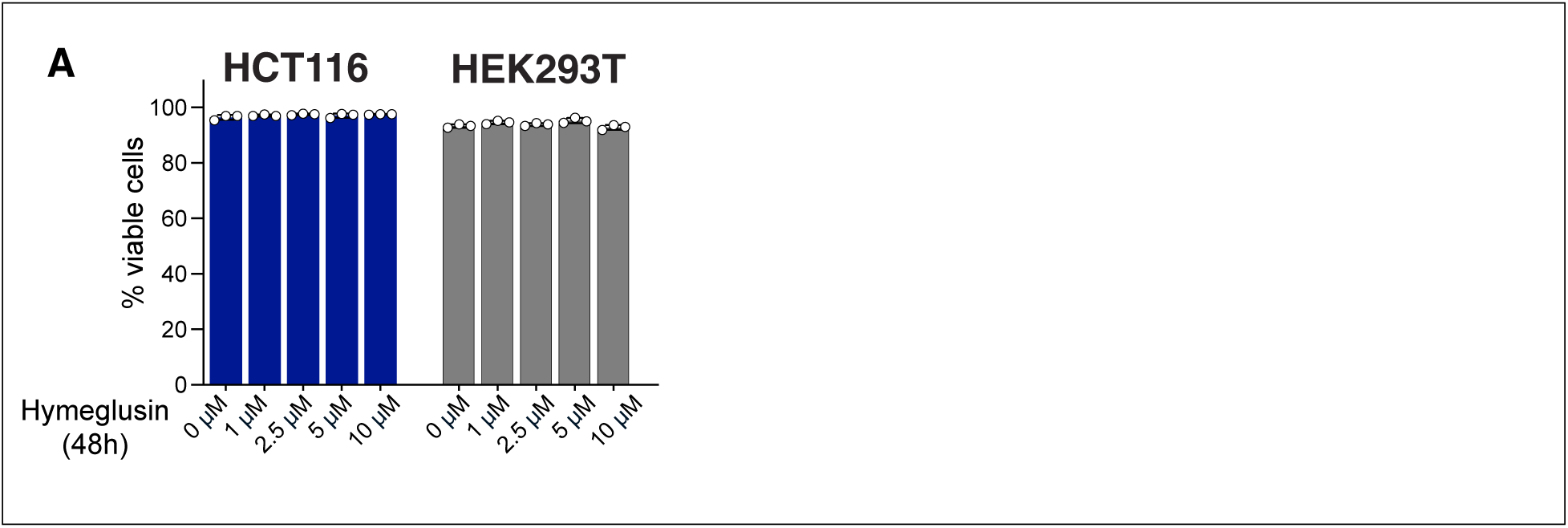
Limited efficacy of Hymeglusin on cell proliferation (Related to Figure 6). (A) HEK293T and HCT116 cells were treated with increasing concentration of Hymeglusin, followed by cell viability assay using flow-cytometry method.

**TABLE S1. AFFINITY-PROTEOMICS USING HYMEGLUSIN-BIOTIN PROBE (RELATED TO FIGURE 2).** HEK293T cells, pretreated with Hymeglusin or DMSO, were further treated with Hymeglusin-biotin, followed by streptavidin enrichment and TMT-6plex analysis.

**TABLE S2. COMPARATIVE PROTEOMICS UPON MVP INHIBITION (RELATED TO FIGURE 3).** TMT-based proteomic data for HCT116 HMGCS1-FKBP12^F36V^ cells following treatment with DMSO, dTAGv1, Hymeglusin, and Simvastatin for 24 hours.

**TABLE S3. KEY RESOURCES TABLE.** Resources of materials used in this study.

